# Genetic diversity and population structure of indigenous chicken in Rwanda using microsatellite markers

**DOI:** 10.1101/825141

**Authors:** R. Habimana, T.O. Okeno, K. Ngeno, S. Mboumba, P. Assami, A. Gbotto, C.T. Keambou, K. Nishimwe, J. Mahoro, N. Yao

## Abstract

Rwanda has about 4.5 million of indigenous chicken (IC) that are very low in productivity. To initiate any genetic improvement programme, IC needs to be accurately characterized. The key purpose of this study was to ascertain the genetic diversity of IC in Rwanda using microsatellite markers. Blood samples of IC sampled from 5 agro-ecological zones were collected from which DNA was extracted, amplified by PCR and genotyped using 28 microsatellite markers. A total of 325 (313 indigenous and 12 exotic) chicken were genotyped and revealed a total number of 305 alleles varying between 2 and 22 with a mean of 10.89 per locus. 186 distinct alleles and 60 private alleles were also observed. The frequency of private alleles was highest in samples from the Eastern region, whereas those from the North West had the lowest. The influx of genes was lower in the Eastern agro-ecological zone than the North West. The mean observed heterozygosity was 0.6155, whereas the average expected heterozygosity was 0.688. The overall inbreeding coefficient among the population was 0.040. Divergence from the Hardy-Weinberg equilibrium was significant in 90% of loci in all the populations. The analysis of molecular variance revealed that about 92% of the total variation originated from variation within populations. Additionally, the study demonstrated that IC in Rwanda could be clustered into four gene groups. In conclusion, there was considerable genetic diversity in IC in Rwanda, which represents a crucial genetic resource that can be conserved or optimized through genetic improvement.

## Introduction

Poultry keeping is an agricultural enterprise with a high potential in Rwanda. More than 40% of households keep poultry out of which approximately 80% consists of indigenous chicken (IC). Raising IC is preferred to exotic breeds because of their small cost of production, scavenging capacity and adaptability to harsh environmental conditions. IC production serves a critical role in the source of revenue for resource-limited countryside families [1]. The productivity of IC in Rwanda, however, is low at an average of 40 to 100 eggs per hen per year and weight ranging from 0.8 to 1.8 kg per year, which is insufficient to meet the needs of the population [2]. This setback has restricted their potential to improve the livelihoods of smallholder farmers thus failing to contribute considerably to the mitigation of poverty in rural areas. To improve the genetic potential of IC in Rwanda, different crossbreeding programmes between IC and exotic chicken were initiated. These programmes, however, are not sustainable because of unpredictable stock and the prohibitive cost of buying and sustaining exotic cocks for breeding purposes in addition to decreased broodiness in the hybridized birds. Additionally, recent global efforts to preserve native genetic resources pose a threat to such programmes, [3]. There is, therefore, the need for an alternative approach to genetic improvement and conservation of IC. Genetic improvement through within-breed selection of IC in Rwanda could be a promising alternative strategy. Nonetheless, genetic enhancements need a resolute breeding objective, sustainable breeding plans, and an in-depth comprehension of the genetic diversity of prevailing genotypes and ecotypes [4]. Therefore, elucidating the genetic characteristics of the prevailing IC stock will not only augment genetic enhancement but will also expedite their preservation [3]. However, there is a scarcity of data on the genetic diversity of IC in Rwanda. The availability of such knowledge could give a clue of the origin and genetic variability in the population and guide selection decisions. As a result, it would be possible to develop apposite mating plans to uphold genetic variation and minimize inbreeding in the population, which would promote response to selection. The current study evaluated the degree of genetic diversity and phylogenetic relationships between populations of IC in Rwanda using simple sequence repeats (SSR) markers.

## Materials and methods

### 2.1 Collection of samples and DNA extraction

In total, 313 distinct IC were sampled from five agro-ecological zones [51, 52, 53, 55, and 102 from Central South (CS), North West (NW), Central North (CN), South West (SW), and East (E), respectively]. Twelve (12) exotic chicken (layers and broilers) were included as a reference for comparison. Populations were reckoned according to agro-ecological zones [5]. A single blood drop was drawn from veins in the wing of each bird and placed on Whatman FTA™ filter cards, left to dry in a cool place for approximately one hour, and held in reserve in discrete envelopes at room temperature awaiting further processing. The isolation of genomic DNA was done using Smith and Burgoyne’s boiling method [6]. The quality of genomic DNA was ascertained through gel electrophoresis using 1% agarose. A NanoDrop Spectrophotometer (Thermo Scientific ™ Nanodrop 2000) was used to quantify the total DNA, which was adjusted to 10ng/μl before use in the subsequent steps of polymerase chain reaction (PCR) and genotyping.

### 2.2 PCR amplification and DNA polymorphism

Twenty-eight fluorescently-labelled polymorphic SSR markers were chosen based on the extent of polymorphism shown by a high polymorphism information content and the genome coverage consistent across previous studies [7]. The PCR reactions had a total volume of 10µl consisting of 30ng target DNA, 5µl of One Taq 2MM and 0.2µl of each forward and reverse primer. The amplifications were done in a thermocycler (Applied Biosystems 9700 Thermal Cycler Gene Amp®) and entailed the first denaturation at 94°C for 3 minutes, 30 cycles of denaturation at 94°C for 30 seconds, the primer annealing at temperatures ranging between 58°C and 64°C based on the primer components (Table 1) for 1 minute, and extension at 72°C for 2 minutes. The last extension step was done at 72°C for 10 minutes. The PCR products of different fluorescent tags were combined according to the exhibited colour and intensity of bands to create uniform signal strength. Hi-Di formimide was used to denature the combined amplicons at 95°C for 3 minutes, this step was followed by capillary electrophoresis separation in an ABI3730 DNA genetic analyzer by using GeneScan-500 Internal LIZ and 1200 Internal LIZ Size Standards. The resultant fragment analysis data and sizes of alleles were counted using GENEMAPPER V 4.1 software (Applied Biosystems).

**Table 1.**
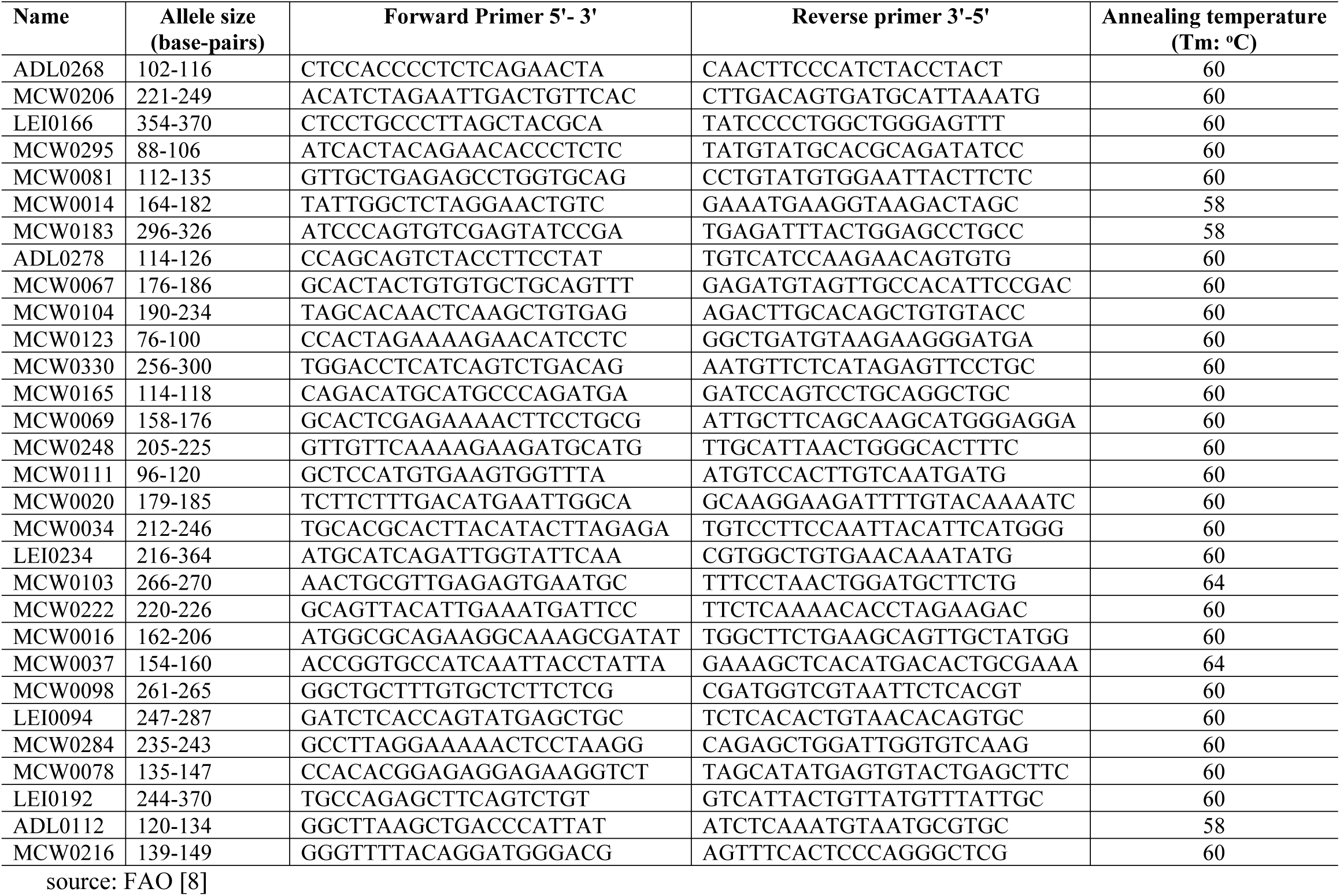
Sequences and physical information of 28 SSR markers used for PCR amplification.

### 2.3 Statistical analysis

#### Genetic diversity and relationship

The polymorphism information content (PIC) was estimated using Powermarker v.3.25 [9]. GenAlEx v.6.5 was used to estimate the allele frequencies, total alleles, expected heterozygosity (He), observed heterozygosity (Ho), and Wright’s F-statistics as well as other parameters such as inbreeding coefficient over all populations (Fis), among populations (Fit) and within populations (Fst) for 28 microsatellite markers [10]. Jackknifing across populations using FSTAT produced standard deviation values that were used to obtain tests of significance per microsatellite locus by creating confidence intervals at 95% and 99% [11].

GENETIX 4.05.2 was used to estimate genetic variation per breed (He, Ho) and the average number of alleles [12]. Gene flow [13] was calculated using Powermarker v.3.25 [9]. Pairwise Fst values, which are indications of the fraction of genetic variation attributed to population sub-structuring, were calculated for various population pairs using GenAlEx v.6.5 [10]. Molecular analysis of variance (AMOVA) was computed using GenAlEx v.6.5 for within and among pre-grouped populations [10]. Powermarker v 3.25 was used to assess genotype frequencies for nonconformity with Hardy-Weinberg equilibrium (HWE) in addition to linkage disequilibrium. GenAlEx v.6.5 [10] was used to approximate Nei’s standard genetic distances [14] among population pairs. The Neighbour-Joining (NJ) programme was used to develop an unrooted NJ cladogram using the Darwin software (v.6.0) according to pairwise kinship distance matrix between populations [15]. A consensus tree assessed by 1,000 bootstraps all through the group of loci was created.

#### Population structure

The possible sum of clusters was approximated using the Evanno method [16] as reported by Dent Earl and Bridgett [17]. A set of rules applied in STRUCTURE was used to group entities based on multi-locus genotypes [18]. The evaluation entailed an admixture model alongside interrelated allele frequencies. During the STRUCTURE analysis, 5 replications of K (presumed sum of subpopulations), extending from 1 to 20 were used together with 100,000 reiterations of Markov Chain Monte Carlo (MCMC) and 50,000 burn-in period in the admixture model. Each estimation of K was redone 5 times to ensure the reproducibility of the outcomes. CLUMPAK (CLUMPAK server), which is a tool used to single out clustering types and bundle population structure deductions across K was used. The Factorial Correspondence Analysis (FCA), which is a multivariate model of analysis, was conducted to observe the associations between entities from unlike zones and to evaluate probable admixtures between the populations. The main variables were the frequencies of alleles at all loci in the populations. The FCA was computed using GENETIX programme [12].

## Results

### 3.1 Genetic diversity

#### Marker Polymorphism across the studied IC populations

The parameters of the variability of the investigated loci are shown in Table 2. Overall, 305 alleles were noted at 28 microsatellite loci with an average of 10.89 alleles per microsatellite marker. The total sum of alleles ranged from 2 (MCW0037) to 22 (LEI0192). The effective number of alleles (NE) ranged between 1.6504 (MCW0078) and 8.901 (LEI0234), with an overall mean of 3.8194. The PIC ranged from 0.3488 (MCW0103) to 0.8775 (LEI0234). Out of the total number of alleles, 20% were private alleles (60), whereas ADL0112 revealed the maximum sum of private alleles (6). The within-population insufficiency in heterozygosity as determined by F_IS_ factor, extended between −1.00 (MCW0037) and 0.338 (LEI0234) with a mean of 0.041 for all loci. The inbreeding coefficient among populations (F_IT_) values ranged from −1.00 (MCW0037) to 0.354 (LEI0234), with a mean of 0.089. Global population differentiation evaluated by F_ST_ was estimated at 0.054. The contribution of 28 microsatellites for population segregation was determined by F_ST_ statistics. F_ST_ values varied from 0.000 (MCW0037) to 0.158 (ADL0268). The overall F-statistics differed significantly from zero (p < 0.05). This differentiation had a significant contribution from all loci. The values for Ho ranged from 0.3015 (MCW0165) to 1 (MCW0037), with an overall mean of 0.6155, while the values of He ranged from 0.394 (MCW0078) to 0.8877 (LEI0234), with a general mean of 0.688. The average number of migrants per generation (Nm) in the whole population and across all the loci was found to be 6.06. About 10% of the loci in all IC populations, did not differ considerably from the HWE.

**Table 2.**
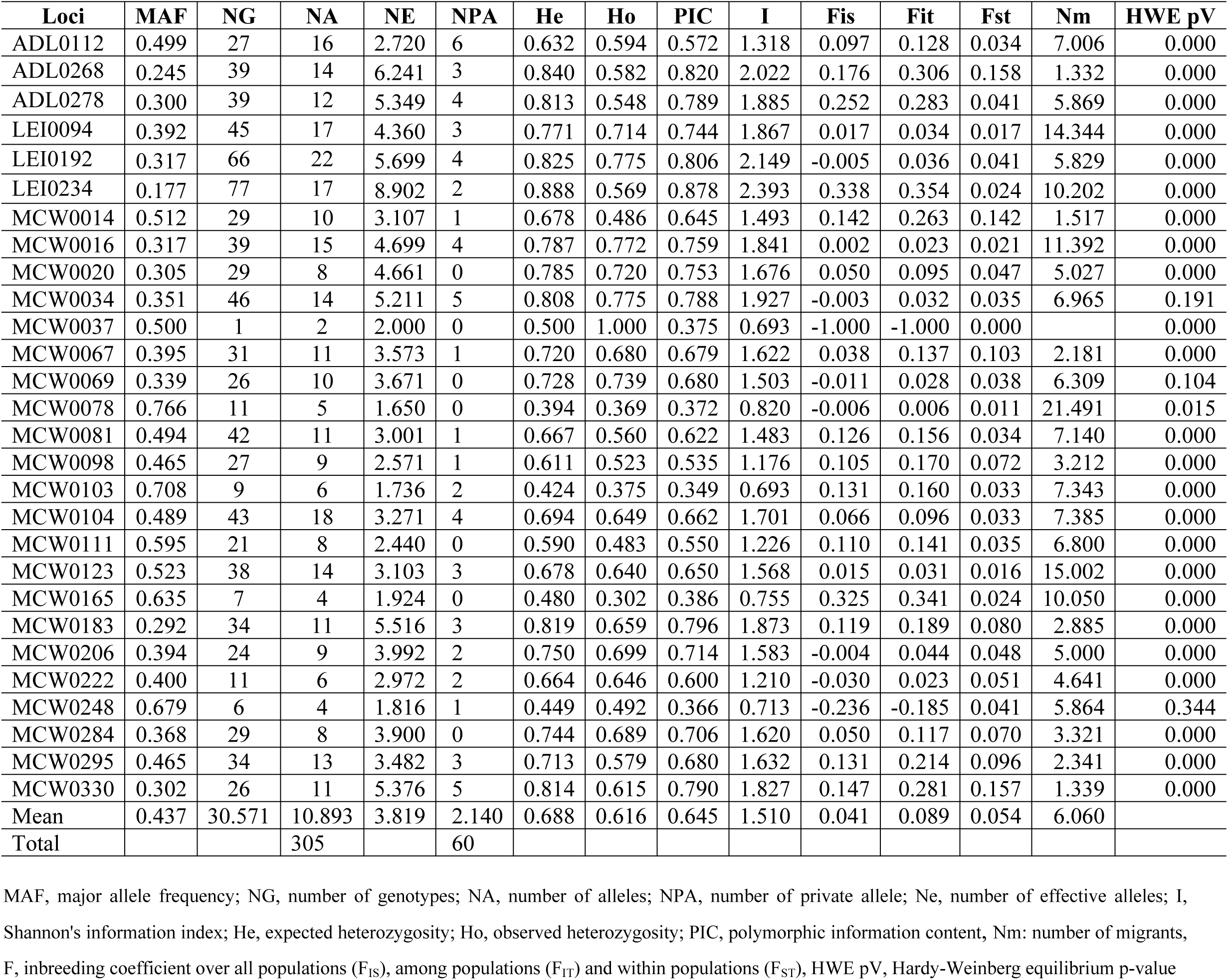
Marker Polymorphism and diversity parameters across studied IC populations in Rwanda.

#### Genetic diversity indices for IC populations from each agro-ecological zone

Genetic diversity indices for IC from each zone is summarized in Table 3. All the loci were polymorphic. The observed frequencies of heterozygote were statistically similar to the expected one (p>0.05), hence, the inbreeding coefficient (F) estimates observed were not substantially different from zero. The mean sum of alleles varied from 5.143 to 8.25. The highest count of alleles (8.2) was found in the Eastern IC population. The highest count of private alleles (21) was observed in the Eastern population, while the NW population did not harbor any private allele. The effective sum of alleles ranged from 3.311 to 3.62. The Shannon Index (I), which is an expression of population diversity in a particular habitat, was high in the SW (1.458) and low in exotic chicken (1.305). Furthermore, the lowest observed heterozygosity was in the CS (0.598) while the highest was recorded in exotic chicken (control) population (0.667). The expected heterozygosity in the populations ranged from0.644 (CN) to 0.680 (SW).

**Table 3.** Common genetic diversity indices as revealed among IC populations in Rwanda.

| Populations | N | %PL | NA | PA | Ne | Ho | He | uHe | F | I |
| --- | --- | --- | --- | --- | --- | --- | --- | --- | --- | --- |
| Central North | 51 | 100 | 6.929 | 6 | 3.354 | 0.623 | 0.644 | 0.650 | 0.021 | 1.322 |
| Central South | 55 | 100 | 7.286 | 15 | 3.359 | 0.598 | 0.661 | 0.668 | 0.077 | 1.372 |
| Exotic chicken | 12 | 100 | 5.143 | 4 | 3.386 | 0.667 | 0.665 | 0.669 | -0.019 | 1.305 |
| East | 102 | 100 | 8.250 | 21 | 3.367 | 0.611 | 0.654 | 0.657 | 0.056 | 1.358 |
| North West | 52 | 100 | 6.500 | 0 | 3.311 | 0.613 | 0.645 | 0.651 | 0.042 | 1.306 |
| South West | 53 | 100 | 7.964 | 14 | 3.620 | 0.626 | 0.680 | 0.686 | 0.063 | 1.458 |
| Total | 325 | 100 | 7.011 | 60 | 3.400 | 0.623 | 0.658 | 0.668 | 0.040 | 1.353 |

NA, number of alleles; PA, number of private allele; Ne, number of effective alleles He, expected heterozygosity Ho, observed heterozygosity uHe: unbiased expected heterozygosity F, inbreeding coefficient I, Shannon's information index.

The p-values of HWE are summarized in Table 4 and confirm that Ho and He do not differ significantly. Thus, taking all the loci into account none of the IC populations diverged from the HWE law.

**Table 4:** Tests for the Hardy-Weinberg equilibrium probability of loci in the IC population in Rwanda.

| Locus | North West | Central North | Central South | East | North-south | Exotic chicken |
| --- | --- | --- | --- | --- | --- | --- |
| ADL0112 | 0.551<sup>ns</sup> | 0.000*** | 0.000*** | 0.003** | 0.000*** | 0.028* |
| ADL0268 | 0.000*** | 0.000*** | 0.163<sup>ns</sup> | 0.000*** | 0.000*** | 0.330<sup>ns</sup> |
| ADL0278 | 0.000*** | 0.000*** | 0.000*** | 0.000*** | 0.003** | 0.349<sup>ns</sup> |
| LEI0094 | 0.001** | 0.976<sup>ns</sup> | 0.000*** | 0.051<sup>ns</sup> | 0.001*** | 0.812<sup>ns</sup> |
| LEI0192 | 0.000*** | 0.000*** | 0.000*** | 0.002** | 0.024* | 0.913<sup>ns</sup> |
| LEI0234 | 0.000*** | 0.000*** | 0.000*** | 0.099<sup>ns</sup> | 0.000*** | 0.720<sup>ns</sup> |
| MCW0014 | 0.000*** | 0.000*** | 0.000*** | 0.000*** | 0.000*** | 0.634<sup>ns</sup> |
| MCW0016 | 0.012* | 0.000*** | 0.000*** | 0.239<sup>ns</sup> | 0.108<sup>ns</sup> | 0.200<sup>ns</sup> |
| MCW0020 | 0.048* | 0.586<sup>ns</sup> | 0.190<sup>ns</sup> | 0.620<sup>ns</sup> | 0.000*** | 0.980<sup>ns</sup> |
| MCW0034 | 0.050* | 0.735<sup>ns</sup> | 0.316<sup>ns</sup> | 0.000*** | 0.816<sup>ns</sup> | 0.412<sup>ns</sup> |
| MCW0037 | 0.000*** | 0.000*** | 0.000*** | 0.000*** | 0.000*** | 0.001*** |
| MCW0067 | 0.000*** | 0.000*** | 0.870<sup>ns</sup> | 0.000*** | 0.000*** | 0.095<sup>ns</sup> |
| MCW0069 | 0.965<sup>ns</sup> | 0.529<sup>ns</sup> | 0.971<sup>ns</sup> | 0.967<sup>ns</sup> | 0.295<sup>ns</sup> | 0.279<sup>ns</sup> |
| MCW0078 | 0.911<sup>ns</sup> | 0.251<sup>ns</sup> | 0.985<sup>ns</sup> | 0.232<sup>ns</sup> | 0.003** | 0.916<sup>ns</sup> |
| MCW0081 | 0.739<sup>ns</sup> | 0.000*** | 0.000*** | 0.000*** | 0.000*** | 0.004** |
| MCW0098 | 0.681<sup>ns</sup> | 0.000*** | 0.000*** | 0.000*** | 0.000*** | 0.005** |
| MCW0103 | 0.012* | 0.752<sup>ns</sup> | 0.000*** | 0.913<sup>ns</sup> | 0.000*** | 0.574<sup>ns</sup> |
| MCW0104 | 0.001** | 1.000<sup>ns</sup> | 0.355<sup>ns</sup> | 0.000*** | 0.755<sup>ns</sup> | 0.213<sup>ns</sup> |
| MCW0111 | 0.046* | 0.189<sup>ns</sup> | 0.127<sup>ns</sup> | 0.003** | 0.687<sup>ns</sup> | 0.545<sup>ns</sup> |
| MCW0123 | 0.503<sup>ns</sup> | 0.909<sup>ns</sup> | 0.000*** | 0.002** | 0.000*** | 0.003** |
| MCW0165 | 0.540<sup>ns</sup> | 0.000*** | 0.004** | 0.000*** | 0.018* | 0.327<sup>ns</sup> |
| MCW0183 | 0.000*** | 0.010* | 0.000*** | 0.000*** | 0.012* | 0.001** |
| MCW0206 | 0.590<sup>ns</sup> | 0.020* | 0.009** | 0.908<sup>ns</sup> | 0.000*** | 0.658<sup>ns</sup> |
| MCW0222 | 0.000*** | 0.096<sup>ns</sup> | 0.000*** | 0.783<sup>ns</sup> | 0.968<sup>ns</sup> | 0.283<sup>ns</sup> |
| MCW0248 | 0.429<sup>ns</sup> | 0.922<sup>ns</sup> | 0.057<sup>ns</sup> | 0.991<sup>ns</sup> | 0.247<sup>ns</sup> | 0.035* |
| MCW0284 | 0.121<sup>ns</sup> | 0.021* | 0.846<sup>ns</sup> | 0.000*** | 0.000*** | 0.437<sup>ns</sup> |
| MCW0295 | 0.279<sup>ns</sup> | 0.000*** | 0.017* | 0.000*** | 0.046* | 0.015* |
| MCW0330 | 0.633<sup>ns</sup> | 0.992<sup>ns</sup> | 0.000*** | 0.000*** | 0.150<sup>ns</sup> | 0.001*** |

ns, not significant, \* P<0.05, \*\* P<0.01, \*\*\* P<0.001

Analysis of molecular variance (AMOVA) revealed that ninety-two percent (92%) of the total variation originated from variation within populations (Table 5).

**Table 5.** Analysis of molecular variance of all loci for the IC population in Rwanda.

| Source | Degree of freedom | Sum square | Mean square | Estimated variances | % of estimated variances |
| --- | --- | --- | --- | --- | --- |
| Among Populations | 5 | 574.201 | 114.840 | 1.838 | 8% |
| Within Populations | 319 | 6346.643 | 19.895 | 19.895 | 92% |
| Total | 324 | 6920.843 |  | 21.733 | 100% |

### 3.2 Genetic relationship

The matrix of pairwise genetic distances between populations (Table 6 and Fig 1) showed a low genetic distance (0.029) between NW and CN populations. A similar trend was observed in SW and CS (0.048). On the other hand, by considering only the IC populations, the highest genetic distance was observed between E and SW populations (0.125). The genetic distance between the IC population in Rwanda and exotic chicken was relatively high (0.231).

**Table 6.** Genetic distance among the IC population in Rwanda.

| Populations | North West | Central North | Central South | Exotic chicken | East |
| --- | --- | --- | --- | --- | --- |
| Central North | 0.029 |  |  |  |  |
| Central South | 0.094 | 0.077 |  |  |  |
| Exotic chicken | 0.199 | 0.213 | 0.231 |  |  |
| East | 0.112 | 0.097 | 0.117 | 0.196 |  |
| South West | 0.104 | 0.092 | 0.048 | 0.118 | 0.125 |

**Fig 1:**
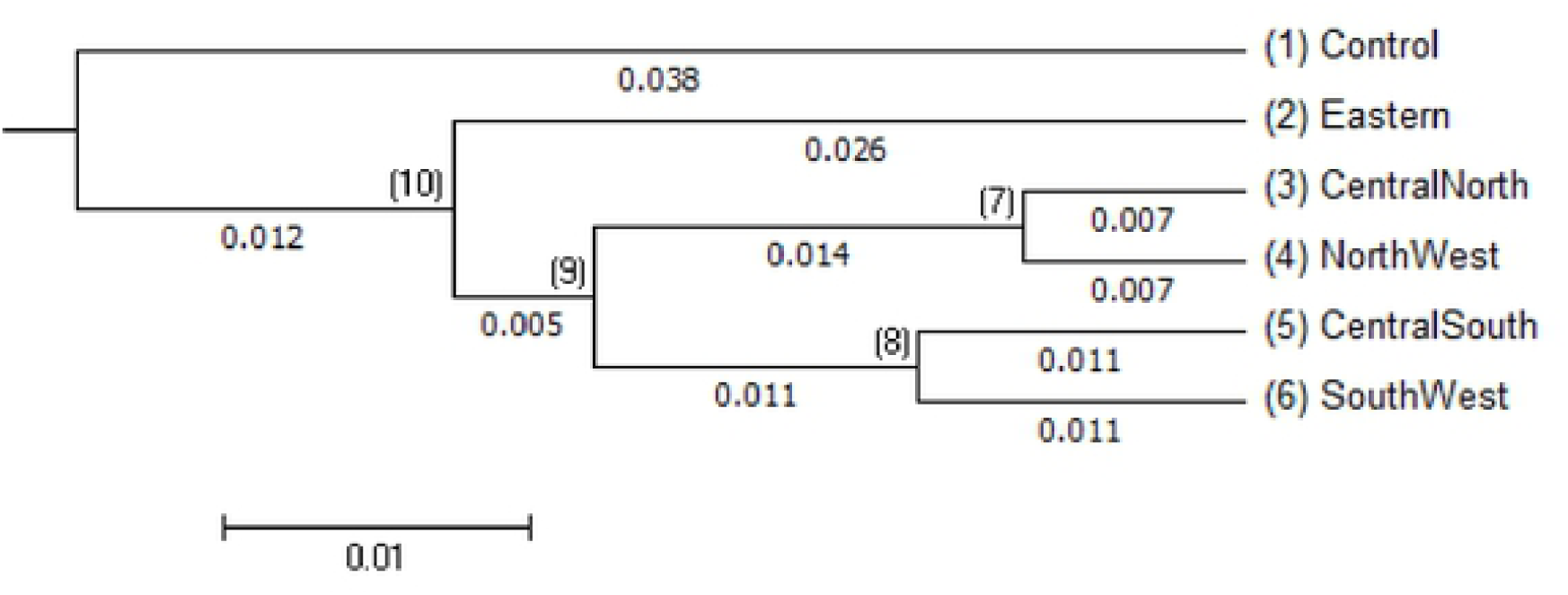
Neighbour-Joining Pair-wise of the IC population in Rwanda.

The extent of genetic distinction among the population with regard to allele frequencies (F_ST_) and gene flow (Nm) are presented in Table 7. The results revealed a low genetic differentiation and a high gene flow between CN and NW, and likewise between SW and CS. A relatively high gene differentiation, however, was found between the E population and other populations.

**Table 7:** Gene flow (upper diagonal) and Gene differentiation (lower diagonal)

| **Populations** | **Central North** | **Central South** | **Exotic chicken** | **East** | **North West** | **South West** |
| --- | --- | --- | --- | --- | --- | --- |
| Central North |  | 2.304 | 1.412 | 2.051 | 6.274 | 2.040 |
| Central South | 0.022 |  | 0.925 | 1.471 | 1.533 | 3.847 |
| Exotic chicken | 0.052 | 0.058 |  | 3.432 | 1.188 | 2.791 |
| East | 0.025 | 0.027 | 0.050 |  | 1.783 | 1.560 |
| North West | 0.012 | 0.026 | 0.053 | 0.028 |  | 1.471 |
| South West | 0.026 | 0.014 | 0.036 | 0.028 | 0.027 |  |

The phylogenetic relationship by the Neighbour-Joining tree showed four (4) IC genetic clusters, namely I, II, III and IV (Fig. 2). The eastern population stands alone unlike the other populations: IC populations from the NW clustered together with those from the CN. Few individuals from the SW population clustered together with the exotic chicken in group III, and finally the rest of SW individuals clustered with those from the CS in group II (Fig 2).

**Fig. 2:**
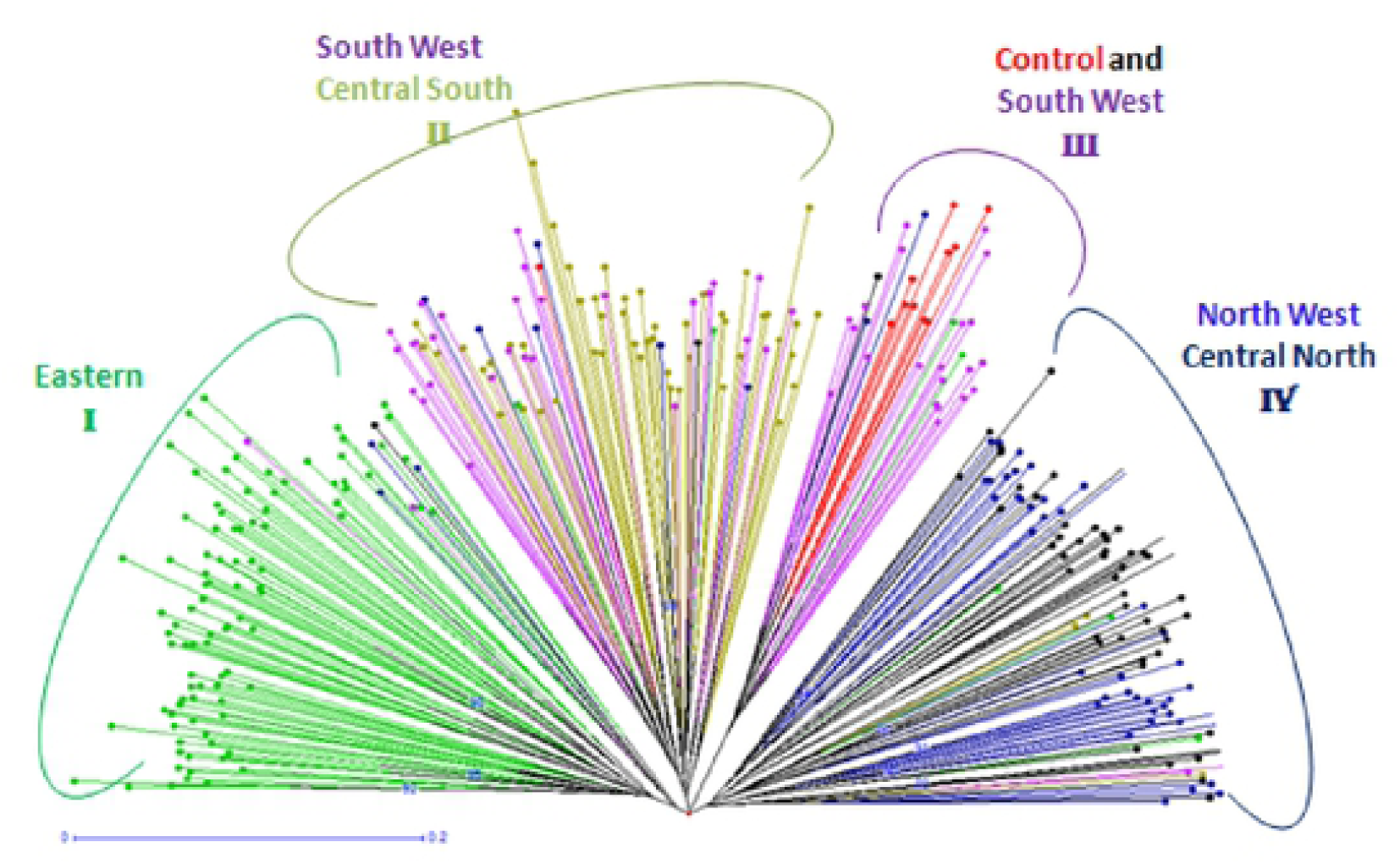
Neighbour-Joining tree of the clustering pattern among IC populations in Rwanda.

### 3.3 Population structure

Data from the Bayesian cluster analysis showed the existence of four (4) main gene pools in the whole IC population in Rwanda. The highest value for *ΔK* was obtained for K = 4 (Table 8 and Fig 3). The first gene pool (I) was composed of CN and NW populations. The second gene pool (II) was made of the Eastern population only. The third (III) included individual from SW and CS and the fourth gene pool (IV) was composed of the remaining individuals of SW and exotic chicken. A high proportion of the admixture was observed in the gene pool III.

**Table 8:** Determination of the number of clusters (K) based on the progression of the average estimate of Ln likelihood of data in IC populations in Rwanda

| K | Reps | Mean LnP(K) | Stddev LnP(K) | Ln'(K) | \|Ln''(K)\| | Delta K |
| --- | --- | --- | --- | --- | --- | --- |
| 1 | 5 | -27680.120000 | 0.192354 | — | — | — |
| 2 | 5 | -26645.700000 | 81.765916 | 1034.420000 | 301.520000 | 3.687600 |
| 3 | 5 | -25912.800000 | 30.968694 | 732.900000 | 82.920000 | 2.677543 |
| 4 | 5 | -25262.820000 | 3.056469 | 649.980000 | 558.300000 | 182.661785 |
| 5 | 5 | -25171.140000 | 37.017671 | 91.680000 | 21.920000 | 0.592150 |
| 6 | 5 | -25057.540000 | 46.761341 | 113.600000 | 19.200000 | 0.410596 |
| 7 | 5 | -24963.140000 | 9.161496 | 94.400000 | 81.200000 | 8.863182 |
| 8 | 5 | -24949.940000 | 63.605566 | 13.200000 | 55.340000 | 0.870050 |
| 9 | 5 | -24881.400000 | 42.680968 | 68.540000 | 29.880000 | 0.700078 |
| 10 | 5 | -24842.740000 | 77.738491 | 38.660000 | 87.640000 | 1.127369 |
| 11 | 5 | -24891.720000 | 114.353824 | -48.980000 | 14.060000 | 0.122952 |
| 12 | 5 | -24954.760000 | 210.975195 | -63.040000 | 330.240000 | 1.565302 |
| 13 | 5 | -24687.560000 | 104.370245 | 267.200000 | 510.500000 | 4.891241 |
| 14 | 5 | -24930.860000 | 402.389690 | -243.300000 | 41.440000 | 0.102985 |
| 15 | 5 | -25132.720000 | 914.525050 | -201.860000 | 542.960000 | 0.593707 |
| 16 | 5 | -24791.620000 | 296.572178 | 341.100000 | 183.320000 | 0.618129 |
| 17 | 5 | -24633.840000 | 54.568333 | 157.780000 | 129.560000 | 2.374271 |
| 18 | 5 | -24605.620000 | 64.775126 | 28.220000 | 204.760000 | 3.161090 |
| 19 | 5 | -24782.160000 | 498.369745 | -176.540000 | 100.700000 | 0.202059 |
| 20 | 5 | -24858.000000 | 559.214181 | -75.840000 | — | — |

**Fig 3:**
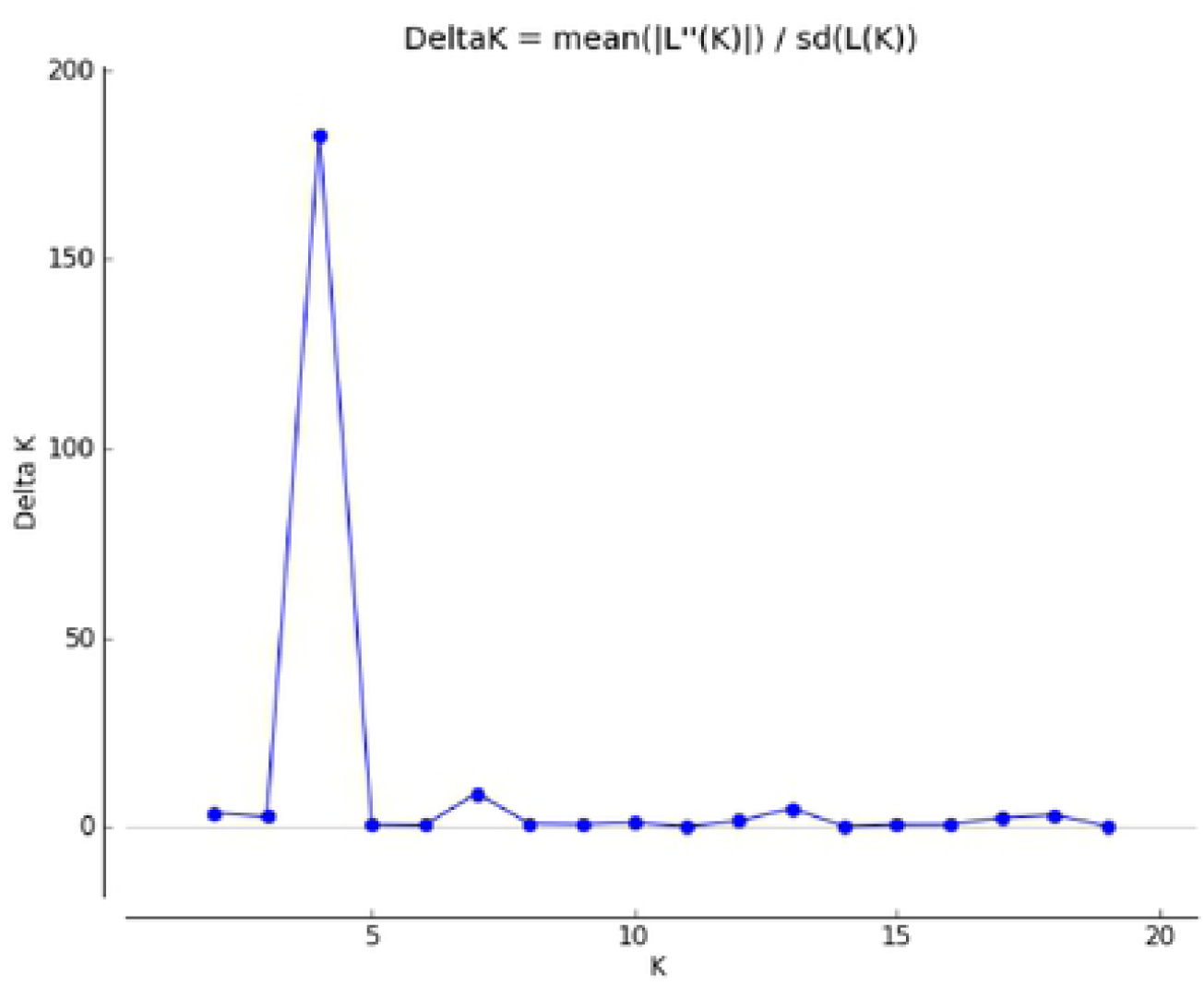
Determination of ΔK approximating the more possible number of clusters in IC populations In Rwanda.

The results of the Factorial Correspondence Analysis (FCA) are shown in Fig 4. It showed tree clusters whereby the Eastern region was still standing alone. NW and CN populations clustered together. Finally, the majority of individuals from the CS, SW and exotic chicken were in the same group.

**Fig 4.**
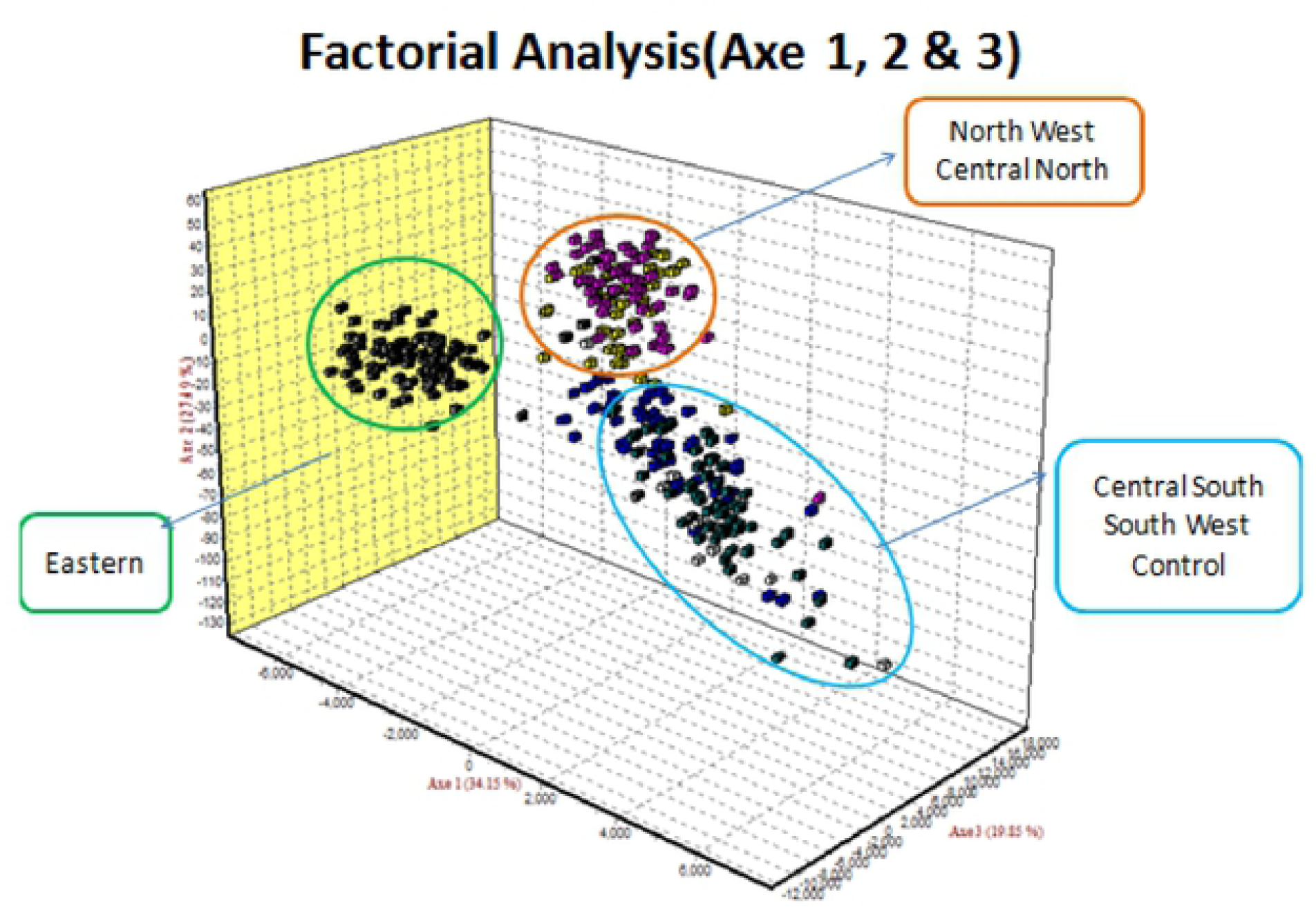
Factorial correspondence analysis.

## Discussion

### 4.1 Genetic diversity

The average PIC was the best index to estimate the polymorphism of alleles [19]. It showed that more information could be obtained from the loci when PIC>0.5. On the other hand, 0.25<PIC<0.5 was an indication of a moderately instructive locus, whereas PIC<0.25 indicated a vaguely informative locus [20]. In this study, 82.3% of all loci were highly informative, which verified that they were suitable for estimating the genetic diversity of IC populations in Rwanda. The highest value of PIC (0.87) was that of LEI0234 and the mean PIC was 0.6451. The PIC values found in this study exceeded those (0.29-080) of Cameroon’s IC [21], and (0.31-0.49) of Chinese IC [21,22], but lower than those obtained by Tang for black-bone IC breeds (0.67) [24]. The mean frequency of alleles per marker found in this study (10.89) exceeded those recorded in previous reports in Cameroon (9.04) [21], in Ghana (7.8) [25], in Iran (5.4) [26], in China (3.8) [27], in Egypt (7.3) [28], in Pakistan (9.1) [29] and in Vietnam (6.41) [30]. The values obtained in this study were, however, lower than those from Brazilian (13.3) [31] and in the same range as from Ethiopian chicken ecotypes (10.6) [32].

The mean number of effective alleles (3.81) obtained in the current study was higher than 3,13 observed in Cameroon [21] and Indian chicken [33]. Heterozygosity can also be considered in genetic diversity. The degree of mean population heterozygosity is an indication of the level of population constancy. Low population heterozygosity informs high population genetic constancy [34]. The present study indicated that Ho of the different IC population varied from 0.3015 to 1 with an overall mean value of 0.6155, while He ranged from 0.394 to 0.887 with an overall average of 0.688.

This study also discovered that the values of Ho and He were similar. As a result, there was no significant difference between zero and the resultant F estimates (0.040), which suggested that the IC populations were in HWE. An implication of this supposition is that the population is under artificial selection, which is indicative of population stability. However, the little variation observed between Ho and He could be attributed to discrepancies in sample size, location, population composition, and the origin of microsatellite markers [35].

The IC populations in Rwanda had a similar level of diversity as their Ethiopian [36], Egyptian [28] and Cameroonian [22] counterparts, but had lower and higher diversity than those observed in southern China [23] and European and Asian IC breeds [25], respectively.

Among Rwanda IC, all populations showed a significantly high degree of inbreeding, which could have an impact on trait fixation in the populations. This degree of inbreeding exceeded that observed for Yunnan IC breeds (0.25) [22] and Turkish IC (0.301) with 10 SSR loci [35].

F_ST_ value (0.054) revealing the diversity between IC populations in Rwanda was higher than 0.048 for Ethiopian IC ecotypes [37] and (0.003-0.040) for Kenyan IC [38] and lower than 0.080 found in Cameroonian IC [21].

### 4.2 Genetic relationships

Wright’s F-statistics strictures showing the inbreeding coefficient in this study was 0.041, which was higher than 0.03 found in Cameroon [21], but was similar to values obtained in many Chinese IC [24, 27]. The F_ST_ permits the approximation of migratory entities in a population per generation (Nm) based on loci. In IC populations in Rwanda, Nm varied from 1.332 to 21.491, with an average of 6.060. This value was higher than that obtained in Cameroun [21].

The number of private alleles (PA) distributed all through the ecotypes showed that there was genetic diversity between populations. In this study, the number of PA was higher in the East (21) followed by CS (15) and SW (14). The NW population, however, did not exhibit any private allele (0). Despite, the number of private alleles being a good indicator of population relationship and structure, further studies need to be carried out to identify possible traits that may be controlled by these private alleles. The total number of private alleles in this study (60) was higher than that found in Cameroun [21].

Findings from AMOVA showed the largest portion of the genetic variation in IC populations in Rwanda existed in individuals within the population (92%). A comparable trend was noted in the Ethiopian [31] and Cameroonian [21] IC ecotypes. The quality of the product, cultural uses of chicken, and the ease with which chicken adapts to the environment are the factors that motivate small-scale farmers to rear IC. These factors highlight the importance of within-population diversity as a key incentive in the rearing of IC [39].

Genetic distance within a population is a useful indicator of separation between various sub-populations. The key assumption of Nei’s standard genetic distance is that hereditary dissimilarities are caused by mutations and genetic drift, whereas Reynolds distance assumes that the increase of genetic differences is due to genetic drift only. The genetic distance between IC populations in SW and CS as well as between NW and CN were not significantly different (P>0.05). It was noted that these regions border each other, thereby implying that there is a high likelihood of sharing genetic materials. Another possible explanation is that these regions could be highly favorable to the IC population or IC populations in these regions could be big enough to prevent mutation and genetic drift. The genetic distances reported in this study fluctuated from to 0.213. These values are in the range of those found in Egyptian IC [28] and in Chinese IC populations [40]. They are, however, higher than those observed in Chinese Bian chicken [23].

When estimating genetic differentiation using allele frequency in such scenarios, the genetic variance between populations can be explained by four major forces, namely, selection, mutation, migration, and genetic drift-[35]. Even though mutation plays a critical role in the long term, short-term evolution is mainly influenced by genetic drift in cases where populations segregated by reproduction[41]. IC populations showed segregation by distance and appeared to be at equipoise under the influence of dispersal and genetic drift. There is a high likelihood that these chickens arrived at their current locations earlier than it had been assumed because there was insufficient time for segregation through distance to come into operation. Furthermore, long-distance gene dispersion is not satisfactorily evident to deter genetic deviation. For this, further investigations need to be conducted using more markers, for example, high-density SNP arrays and mitochondrial DNA.

### 4.3 Population structure

The genetic similarity in a collection of breeds with high diversity can be resolved efficiently by cluster analysis, which facilitates the identification of individuals with similar or diverse multi-locus genotypes [42]. In our study, the cluster based on the neighbour-joining approach revealed grouping arrays of association and genetic relationships among individuals. These individuals were grouped in four clusters formed by ecotypes from distinct collection sites (NW and CN; SW1 and CS; SW2 and control and finally, East stands alone). This was confirmed by the STRUCTURE analysis which revealed four gene pools across IC in Rwanda. These gene pools are distributed exactly according to the different clusters as shown by the neighbour-joining method. The observed gene pools could be accounted for by the sum of private alleles recorded in the population besides the genetic distance between populations. For example, the Eastern region recorded the highest frequency of private alleles, whereas the NW had the lowest number. This observation could be attributed to the large population of IC in the Eastern region out of all the study sites, which minimized gene inflow in this area. Conversely, the lowest number of IC was noted in the NW region, which could be interpreted to mean that the majority of people in this area either buy chicken or exchange cocks from the neighbouring areas such as CN. Consequently, there is a high influx of genes in these regions This is not surprising since these areas border each other geographically. These findings corroborated the observations of a study conducted in Kenya where the Mantel test had uncovered a positive association between hereditary and geographic distances [43]. Our study also confirmed that geographic distances affected the population’s genetic structure [43]. The portion of SW chicken populations that clustered with the exotic chicken (control) could be attributed to the fact that different crossing programmes between IC and improved chicken breeds have been introduced in that region to improve the genetic potential of IC in Rwanda [44].

#### Conclusion

The results portrayed by this study are the first to recount the genetic diversity and constitution of IC from Rwanda. Overall, the IC populations in Rwanda had high levels of significant genetic variability as per different genetic diversity parameters applied in this study. Therefore, data on genetic diversity estimated by assimilating within and between population variances may inform preservation strategies and the better establishment of priorities. In addition, this study found that IC in Rwanda belongs to four major gene pools that could be preserved independently to uphold their genetic diversity. Generally, these findings provide the fundamental step in the direction of judicious decision-making before the development of genetic enhancement and preservation programmes without interfering with the uniqueness of IC in Rwanda.

## Acknowledgement

The authors acknowledge the financial and technical support from BecA-ILRI Hub through Africa Biosciences Challenge Fund (ABCF) programmes. The ABCF Programmes is funded by the Australian Department for Foreign Affairs and Trade (DFAT) through the BecA-CSIRO partnership; the Syngenta Foundation for Sustainable Agriculture (SFSA); the Bill & Melinda Gates Foundation (BMGF); the UK Department for International Development (DFID) and the Swedish International Development Cooperation Agency (SIDA).

This material is also based upon work supported by the United States Agency for International Development, as part of the Feed the Future initiative, under the CGIAR Fund, award number BFS-G-11-00002, and the predecessor fund the Food Security and Crisis Mitigation II grant, award number EEM-G-00-04-00013.

## Author Contributions

**Conceptualization:** Richard Habimana, Nasser Yao, Pauline Assami, Tobias Okeno and Kiplangat Ngeno

**Data curation:** Richard Habimana

**Formal analysis:** Richard Habimana, Nasser Yao, Pauline Assami and Anique Gbotto

**Funding acquisition:** Nasser Yao, Richard Habimana, and Pauline Assami

**Investigation:** Richard Habimana, Kizito Nishimwe, Christian Keambou and Janvier Mahoro

**Methodology:** Richard Habimana, Yao Nasser, Christian Keambou and Kizito Nishimwe

**Project administration:** Nasser Yao, Pauline Assami and Richard Habimana.

**Resources:** Richard Habimana, Nasser Yao, Tobias Okeno, Kiplangat Ngeno, Christian Keambou and Pauline Assami.

**Supervision:** Nasser Yao, Tobias Okeno and Kiplangat Ngeno

**Visualization:** Richard Habimana and Sylvere Mboumba

**Writing ± original draft:** Richard Habimana.

**Writing ± review & editing:** Richard Habimana, Nasser Yao, Tobias Okeno and Kiplangat Ngeno

